# EvoAug: improving generalization and interpretability of genomic deep neural networks with evolution-inspired data augmentations

**DOI:** 10.1101/2022.11.03.515117

**Authors:** Nicholas Keone Lee, Ziqi Tang, Shushan Toneyan, Peter K Koo

## Abstract

Deep neural networks (DNNs) hold promise for functional genomics prediction, but their generalization capability may be limited by the amount of available data. To address this, we propose EvoAug, a suite of evolution-inspired augmentations that enhance the training of genomic DNNs by increasing genetic variation. However, random transformation of DNA sequences can potentially alter their function in unknown ways. Thus, we employ a fine-tuning procedure using the original non-transformed data to preserve functional integrity. Our results demonstrate that EvoAug substantially improves the generalization and interpretability of established DNNs across prominent regulatory genomics prediction tasks, offering a robust solution for genomic DNNs.

---

Uncovering *cis*-regulatory elements and their coordinated interactions is a major goal of regulatory genomics. Deep neural networks (DNNs) offer a promising avenue to learn these genomic features *de novo* through being trained to take DNA sequences as input and predict their regulatory functions as output^1–3^. Following training, these DNNs have been employed to score the functional effect of disease-associated variants^4, 5^. Moreover, *post hoc* model interpretability methods have revealed that DNNs base their decisions on learning sequence motifs of transcription factor (TF) binding sites and dependencies with other TFs and sequence context^6–10^.

For DNNs, generalization typically improves with more training data. However, the amount of data generated in a high-throughput functional genomics experiment is fundamentally limited by the underlying biology. For example, the extent to which certain TFs bind to DNA is constrained by the availability of high-affinity binding sites in accessible chromatin.

To expand a finite dataset, data augmentations can provide additional variations on existing training data^11, 12^. Data augmentations act as a form of regularization, guiding the learned function to be invariant to symmetries created by the data transformations^13, 14^. This approach can help prevent a DNN from overfitting to spurious features and improve generalization^15^. The main challenge with data augmentations in genomics is quantifying how the regulatory function changes for a given transformation. With image data, basic affine transformations can translate, magnify, or rotate an image without changing its label. However, in genomics, the available neutral augmentations are reverse-complement transformation^16^ and small random translations of the input sequence^17, 18^. With the finite size of experimental data and a paucity of augmentation methods, strategies to promote generalization for genomic DNNs are limited.

Here we introduce EvoAug, an open-source PyTorch package that provides a suite of evolution-inspired data augmentations. We show that training DNNs with EvoAug leads to better generalization performance and improves efficacy with standard *post hoc* explanation methods, including filter interpretability and attribution analysis, across prominent regulatory genomics prediction tasks for well-established DNNs.

## Results and Discussion

### Evolution-inspired data augmentations for sequence-based genomic DNNs

To enhance the effectiveness of sequence-based models, data augmentations should aim to increase genetic diversity while maintaining the same biological functionality. Evolution provides a natural process to generate genetic variability, including random mutations, deletions, insertions, inversions, and translocations, among others^19^. However, these genetic changes often have functional consequences that expand phenotypic diversity and aid in natural selection. While the addition of homologous sequences to a dataset could achieve the goal of increasing sequence diversity while preserving biological function, identifying regulatory regions with similar functions throughout the genomes across species is difficult. Alternatively, synthetic perturbations that do not alter the function can be applied, but it is crucial to have prior knowledge to ensure that features such as motifs and their dependencies are not affected. Therefore, formulating new data augmentation strategies for genomics remains a significant challenge.

In this study, we present a suite of evolution-based data augmentations and a two-stage training curriculum to preserve functional integrity (Fig. 1a). In the first stage, a DNN is trained on sequences with EvoAug augmentations applied stochastically online during training, using the same training labels as the wild-type sequence. The goal is to enhance the model’s ability to learn robust representations of features, such as motifs, by exposing it to expanded (albeit synthetically generated) genetic variation. While each augmentation has the potential to disrupt core motifs in any given perturbation, we expect the overall effect to preserve motifs on average. However, the specific data augmentations employed may introduce a bias in how these motif grammars are structured. Thus, in the second stage, the DNN is fine-tuned on the original, unperturbed data to refine these features and guide the function towards the observed biology, thereby removing any bias introduced by the data augmentations (see Methods).

**Figure 1.**
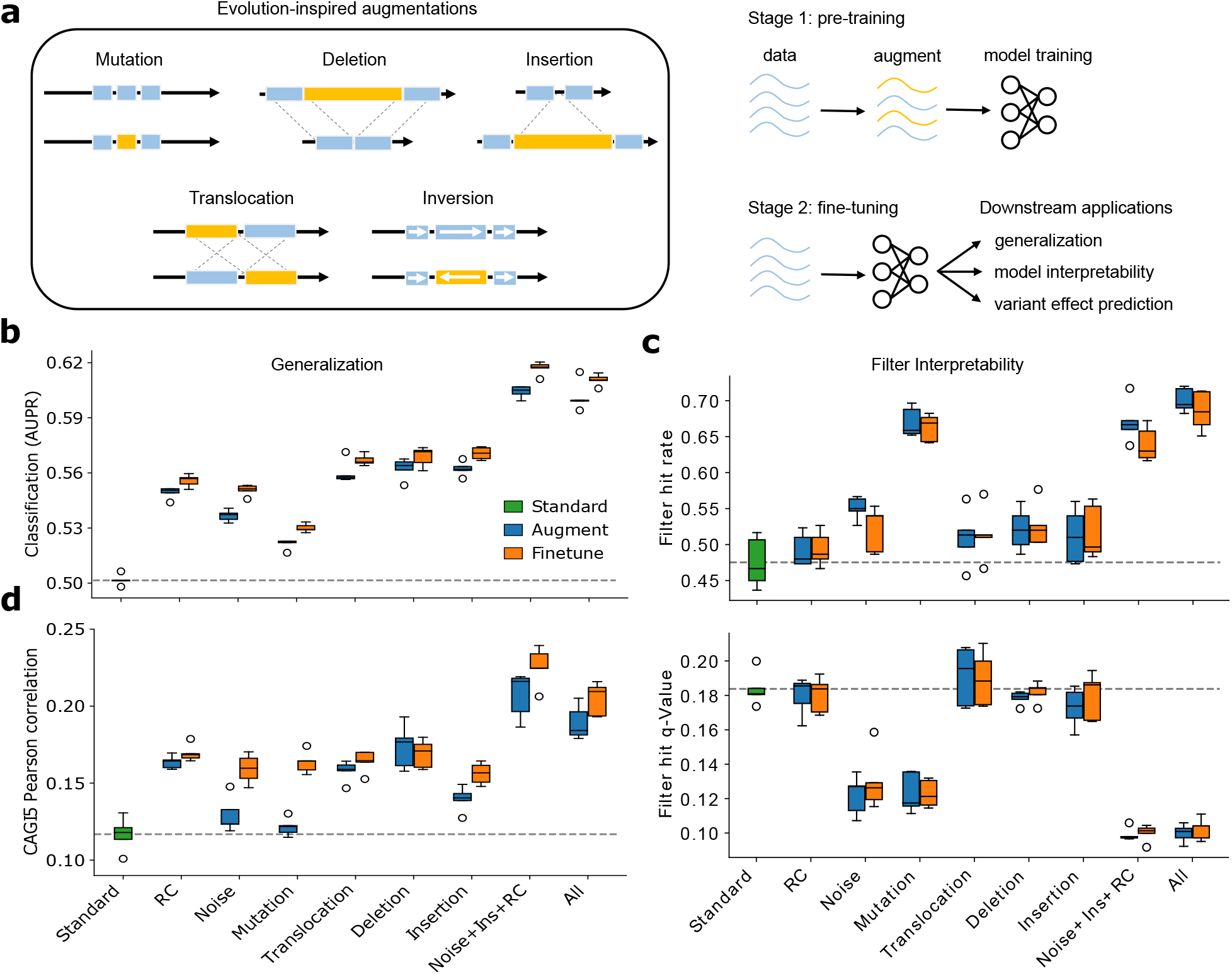
EvoAug improves generalization and interpretability of Basset models. (**a**) Schematic of evolution-inspired data augmentations (left) and the two-stage training curriculum (right). (**b**) Generalization performance (area under the precision-recall curve) for Basset models pretrained with individual and combinations of augmentations, i.e., Noise+Ins+RC (Gaussian noise, insertion, reverse-complement) and all augmentations (Gaussian noise, reverse-complement, mutation, translocation, deletion, insertion), and fine-tuned on Basset dataset. Standard represents no augmentations during training. (**c**) Comparison of the average hit rate of first-layer filters to known motifs in the JASPAR database (top) and the average *q*-value of the filters with matches (bottom). (**d**) Comparison of the average Pearson correlation between model predictions and experimental data from CAGI5 Challenge. (**b**-**d**) Each box-plot represents 5 trials with random initializations.

EvoAug data augmentations introduce a modeling bias to learn invariances of the (un)natural symmetries generated by the augmentations. For instance, random insertions and deletions assume that the distance between motifs is not critical, whereas random inversions and translocations promote invariances to motif strand orientation and the order of motifs, respectively. Nevertheless, the bias created by the augmentations can lead to poor generalization when the introduced bias does not accurately reflect the underlying biology. Therefore, the fine-tuning stage is critical as it provides an avenue to unlearn any biases not supported by the observed data.

### EvoAug improves generalization and interpretability of genomic DNNs

To demonstrate the utility of EvoAug, we analyzed several established DNNs across three prominent types of regulatory genomic prediction tasks that span a range of complexity.

First, we applied Evoaug to the Basset model and dataset^20^, which consists of a multi-task binary classification of chromatin accessibility sites across 161 cell types/tissues. We trained the Basset model with each augmentation applied independently and in various combinations. We conducted a hyperparameter sweep to determine the optimal settings for each augmentation (Supplementary Figs. 1–5). From hyperparameter sweeps, we observed that the inversion augmentation improved performance up to the sequence length, which is essentially a reverse-complement transformation (Supplementary Figs. 1, 3, and 4). Hence, inversions were excluded to reduce redundancy.

Remarkably, EvoAug-trained DNNs outperformed standard training with no augmentations (Fig. 1b). The best results were achieved when multiple augmentations were used together. Additionally, we found that fine-tuning on the original data further improved performance, even when augmentation hyperparameters were poorly specified (Supplementary Fig. 1). Notably, specific EvoAug augmentations, such as random mutations and combinations of data augmentations, had a profound impact on improving the motif representations learned by the first-layer convolutional filters (Fig. 1c). The convolutional filters capture a wider repertoire of motifs and their representations better reflect known motifs, both quantitatively and qualitatively, when compared with convolutional filters of models trained without augmentations. This suggests that EvoAug augmentations can help DNNs learn more accurate and informative representations of the sequence motifs.

A major downstream application of genomic DNNs is to score the functional consequences of non-coding mutations. By evaluating the zero-shot prediction capabilities of each DNN on saturation mutagenesis data of 15 *cis*-regulatory elements from the CAGI5 Challenge^21^, we found that models trained with EvoAug outperformed their standard training counterpart (Fig. 1d). Notably, Basset’s performance was comparable to other DNNs based on binary predictions^17^; however, its overall performance was lower than more sophisticated DNNs and top competitors in the CAGI5 challenge^2^. Interestingly, we observed that DNNs pretrained with Gaussian noise or random mutagenesis augmentations did not perform well. These augmentations impose flatness locally in sequence-function space, effectively reducing the effect size of nucleotide variants. However, fine-tuning these models improved their variant effect predictions beyond what was achieved with standard training, thus demonstrating the effectiveness of the two-stage training curriculum.

To further demonstrate the benefits of EvoAug, we trained DeepSTARR models as a multi-task quantitative regression to predict enhancer activity from self-transcribing active regulatory region sequencing (STARR-seq) data^9^, where each task represents a different promoter from a developmental or housekeeping gene in *Drosophila* S2 cells. Most EvoAug augmentations resulted in improved performance, except for reverse-complement and random mutations (Fig. 2a and Supplementary Figures 3–5). As before, we observed additional performance gains when augmentations were used in combination. Furthermore, the attribution maps generated by EvoAug-trained models were more interpretable, with identifiable motifs and less spurious noise (Supplementary Fig. 6).

**Figure 2.**
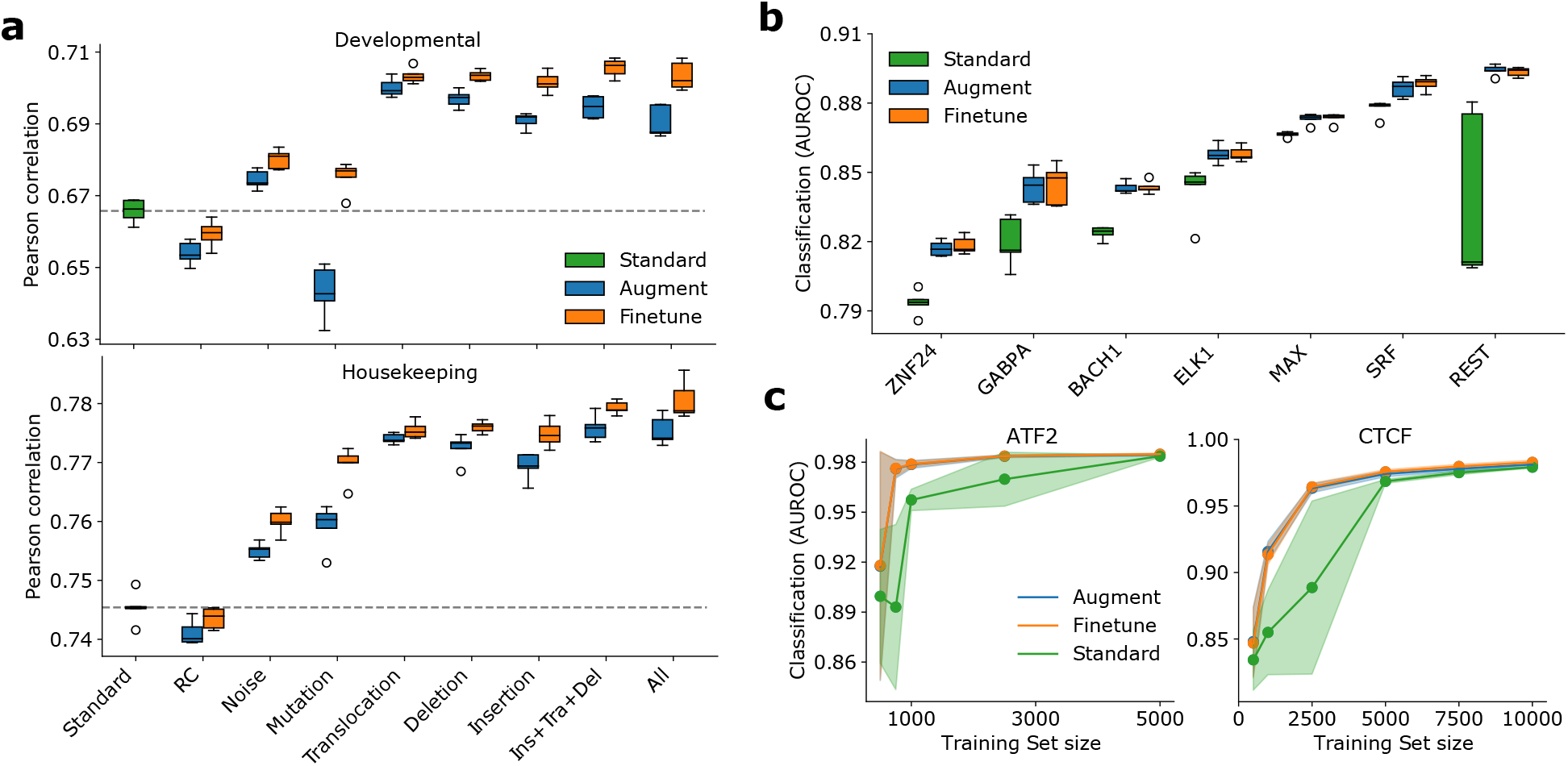
Generalization of EvoAug on additional models and datasets. (**a**) Box-plot of regression performance for DeepSTARR models pretrained with individual or combination of augmentations (i.e. insertion + translocation + deletion; all augmentations) and fine-tuned on original STARR-seq data for two promoters: developmental (top) and housekeeping (bottom). Standard represents no augmentations during training. (**b**) Box-plot of classification performance (area under the receiver-operating-characteristic curve) for DNNs trained on ChIP-seq datas. **(c)**Average classification performance for ChIP-seq experiments downsampled to different dataset sizes. Shaded region represents the standard deviation of the mean. (**a**-**b**) Each box-plot represents 5 trials with random initializations.

In addition, we found that the EvoAug-trained DNNs consistently outperformed DNNs with standard training on various single-task binary classification tasks for TF binding across multiple chromatin immunoprecipitation sequencing (ChIP-seq) datasets (Fig. 2b). Interestingly, we did not observe any significant improvement in performance after fine-tuning, suggesting that the implicit prior imposed by EvoAug augmentations was appropriate for these tasks; the underlying regulatory grammars for these TFs are not complex.

To further investigate the impact of EvoAug on small datasets, we retrained each DNN on down-sampled versions of two abundant ChIP-seq datasets. We found that EvoAug-trained DNNs exhibit a greater improvement in performance for smaller datasets compared to standard training (Fig. 2c). This result suggests that EvoAug can be particularly useful in scenarios where the available training data is limited.

Training with EvoAug adds a computational cost, depending on the augmentations chosen and their settings (Supplementary Tables 2 and 3). Nevertheless, EvoAug stabilized training (Supplementary Fig. 7), leading to smoother convergence and improved generalization overall.

## Conclusion

EvoAug greatly expands the set of available data augmentations for genomic DNNs. Our study demonstrated that EvoAug’s two-stage training curriculum is effective in improving generalization performance. Moreover, EvoAug-trained models learned better representations of consensus motifs, as evidenced by filter visualization and attribution analysis.

Our findings support previous arguments for using evolution as a natural source of data augmentation^22^. Interestingly, the impact of synthetic evolutionary perturbations was not excessively disruptive, and performance even improved before fine-tuning in most cases. This functional robustness appears to be a characteristic of the non-coding genome^23^.

Data augmentations are a commonly used technique to balance bias and variance in machine learning models. However, their effectiveness is expected to decrease as the dataset size increases. Nevertheless, EvoAug still improved performance on the already large Basset dataset. Other methods that can enhance generalization include multitask learning^24^, contrastive learning^25, 26^, and language modeling^27^. Even though Basset and DeepSTARR are already trained in a multitask framework, EvoAug improved their performance. Multitasking can introduce class imbalance, but EvoAug provides additional examples with pseudo-positive labels, which can mitigate this issue. EvoAug also provides different views of the data, which can be useful for contrastive learning. Importantly, EvoAug is a lightweight and effective strategy that only requires the original data.

The optimal combination of augmentations and their hyperparameter choices depends on the model and dataset. While we performed hyperparameter grid searches in this study, more advanced search strategies such as population-based training^28^ using Ray Tune^29^ could improve efficiency. In the future, we plan to investigate EvoAug’s potential in cross-dataset generalization and variant effect predictions, including expression quantitative trait loci.

EvoAug is a PyTorch package that is open-source, easy to use, extensible, and accessible via pip (https://pypi.org/project/evoaug) and GitHub (https://github.com/p-koo/evoaug), with full documentation provided on ReadtheDocs.org (https://evoaug.readthedocs.io.). In time, we plan to extend EvoAug functionality to TensorFlow^30^ and JAX^31^. We anticipate that EvoAug will have broad utility in improving the efficacy of sequence-based DNNs for regulatory genomics.

## Methods

### Models and Datasets

#### Basset

The Basset dataset^20^ consists of a multi-task binary classification of chromatin accessibility sites across 161 cell types/tissues. The inputs are genomic sequences of length 600 nt and the output are binary labels (representing accessible or not accessible) for 161 cell types measured experimentally using DNase I hypersensitive sites sequencing (DNase-seq). We filtered sequences that contained at least one N character and the data splits (training; validation; test) reduced from (1,879,982; 70,000; 71,886) to (437,478; 16,410; 16,703). This “cleaned” dataset was analyzed using a Basset-inspired model, which is given according to:

- Input *x* ∈ {0, 1}^600×4^ (one-hot encoding of 600 nt sequence)
- 1D convolution (300 filters, size 19, stride 1)
- BatchNorm + ReLU
- Max-pooling (size 3, stride 3)
- 1D convolution (200 filters, size 11, stride 1)
- BatchNorm + ReLU
- Max-pooling (size 4, stride 4)
- 1D convolution (200 filters, size 7, stride 1)
- BatchNorm + ReLU
- Max-pooling (size 2, stride 2)
- Fully-connected (1000 units)
- BatchNorm + ReLU
- Dropout (0.3)
- Fully-connected (1000 units)
- BatchNorm + ReLU
- Dropout (0.3)
- Fully-connected output (161 units, sigmoid)

BatchNorm represents batch normalization^32^, and dropout^33^ rates set the probability that neurons in a given layer are temporarily removed during each mini-batch of training.

#### DeepSTARR

The DeepSTARR dataset^9^ consists of a multi-task regression of enhancer activity for two promoters, well-known developmental and housekeeping transcriptional programs in *D. melanogaster* S2 cells. The inputs are genomic sequences of length 249 nt and the output is 2 scalar values representing the activity of developmental enhancers and housekeeping enhancers measured experimentally using STARR-seq. Sequences with N characters were also removed, but this minimally affected the size of the dataset (i.e., reduced it by approximately 0.005%). This dataset was analyzed using the original DeepSTARR model, given according to:

- Input *x* ∈ {0, 1}^249×4^
- 1D convolution (256 filters, size 7, stride 1)
- BatchNorm + ReLU
- Max-pooling (size 2, stride 2)
- 1D convolution (60 filters, size 3, stride 1)
- BatchNorm + ReLU
- Max-pooling (size 2, stride 2)
- 1D convolution (60 filters, size 5, stride 1)
- BatchNorm + ReLU
- Max-pooling (size 2, stride 2)
- 1D convolution (120 filters, size 3, stride 1)
- BatchNorm + ReLU
- Max-pooling (size 2, stride 2)
- Fully-connected (256 units)
- BatchNorm + ReLU
- Dropout (0.4)
- Fully-connected (256 units)
- BatchNorm + ReLU
- Dropout (0.4)
- Fully-connected output (2 units, linear)

#### ChIP-seq

Transcription factor (TF) chromatin immunoprecipitation sequencing (ChIP-seq) data was processed and framed as a binary classification task. The inputs are genomic sequences of length 200 nt and the output is a single binary label representing TF binding activity, with positive-label sequences indicating the presence of a ChIP-seq peak and negative-label sequences indicating a peak for a DNase I hypersensitive site from the same cell type but one that does not overlap with any ChIP-seq peaks. Nine representative TF ChIP-seq experiments in a GM12878 cell line and a DNase-seq experiment for the same cell line were downloaded from ENCODE^34^; for details, see Supplementary Table 1. Negative sequences (i.e., DNase-seq peaks that do not overlap with any positive peaks) were randomly down-sampled to match the number of positive sequences, keeping the classes balanced. The dataset was split randomly into training, validation, and test set according to the fractions 0.7, 0.1, and 0.2, respectively.

A custom convolutional neural network was employed to analyze these datasets, given according to:

- Input *x* ∈ {0, 1}^200×4^
- 1D convolution (64 filters, size 7, stride 1)
- BatchNorm + ReLU
- Dropout (0.2)
- Max-pooling (size 4, stride 4)
- 1D convolution (96 filters, size 5, stride 1)
- BatchNorm + ReLU
- Dropout (0.2)
- Max-pooling (size 4, stride 4)
- 1D convolution (128 filters, size 5, stride 1)
- BatchNorm + ReLU
- Dropout (0.2)
- Max-pooling (size 2, stride 2)
- Fully-connected layer (256 units)
- BatchNorm + ReLU
- Dropout (0.5)
- Fully-connected output layer (1 unit, sigmoid)

### Evolution-inspired Data Augmentations

EvoAug is comprised of a set of data augmentations given by the following:

- Mutation: a transformation where single nucleotide mutations are randomly applied to a given wild-type sequence. This is implemented as follows: (1) given the hyperparameter of the fraction of nucleotides in each sequence to mutate (`mutate_frac`), the number of mutations for a given sequence length is calculated; (2) a position along the sequence is randomly sampled (with replacement) for each number of mutations; and (3) the selected positions are mutagenized to a random nucleotide. Since our implementation does not guarantee that a nucleotide selected will be mutated to a different nucleotide than it originally was, we take approximate account for silent mutations by dividing the user-defined `mutate_frac` by 0.75 so that on average the fraction of nucleotides in each sequence mutated to a different nucleotide is equal to `mutate_frac`.
- Translocation: a transformation that randomly selects a break point in the sequence (thereby creating two segments) and then swaps the order of the two sequence segments. An equivalent statement of this transformation is a “roll”—shifting the sequence forward along its length a randomly specified distance and then reintroducing the part of the sequence shifted beyond the last position back at the first position. This is implemented as follows: (1) given the hyperparameters of the minimum distance (`shift_min`, default 0) and maximum distance (`shift_max`) of the shift, the integer-valued shift length is chosen randomly from the interval `[–shift_max, –shift_min] ∪ [shift_min, shift_max]`, where a negative value simply denotes a backward shift rather than a forward shift; and (2) the shift is applied to the sequence with a `roll()` function in PyTorch.
- Insertion: a transformation where a random DNA sequence (of random length) is inserted randomly into a wild-type sequence. This is implemented as follows: (1) given the hyperparameters of the minimum length (`insert_min`, default 0) and maximum length (`insert_max`) of the insertion, the integer-valued insertion length is chosen randomly from the interval between `insert_min` and `insert_max` (inclusive); and (2) the insertion is inserted at a random position within the original sequence. Importantly, to maintain a constant input sequence length to the model (i.e., original length plus `insert_max`), the remaining amount of length between the insertion length and `insert_max` is split evenly and placed on the 5’ and 3’ flanks of the sequence, with the remainder from odd lengths going to the 3’ end. Whenever an insertion augmentation is employed in combination with other augmentations, all sequences without an insertion are padded with a stretch of random DNA of length `insert_max` at the 3’ end to ensure that the model processes sequences with a constant length for both training and inference time.
- Deletion: a transformation where a random, contiguous segment of a wild-type sequence is removed, and the shortened sequence is then padded with random DNA sequence to maintain the same length as wild-type. This is implemented as follows: (1) given the hyperparameters of the minimum length (`delete_min`, default 0) and maximum length (`delete_max`) of the deletion, the integer-valued deletion length is chosen randomly from the interval between `delete_min` and `delete_max` (inclusive); (2) the starting position of the deletion is chosen randomly from the valid positions in the sequence that can encapsulate the deletion; (3) the deletion is performed on the designated stretch of the sequence; (4) the remaining portions of the sequence are concatenated together; and (5) random DNA is used to pad the 5’ and 3’ flanks to maintain a constant input sequence length, similar to the procedure for insertions.
- Inversion: a transformation where a random subsequence is replaced by its reverse-complement. This is implemented as follows: (1) given the hyperparameters of the minimum length (`invert_min`, default 0) and maximum length (`invert_max`) of the inversion, the integer-valued inversion length is chosen randomly from the interval between `invert_min` and `invert_max` (inclusive); (2) the starting position of the inversion is chosen randomly from the valid position indices in the sequence; and (3) the inversion (i.e., a reverse-complement transformation) is performed on the designated subsequence while the remaining portions of the sequence remain untouched.
- Reverse-complement: a transformation where a full sequence is replaced with some probability `rc_prob` by its reverse-complement.
- Gaussian noise: a transformation where Gaussian noise (with distribution parameters `noise_mean = 0` and `noise_std`) is added to the input sequence; a random value drawn independently and identically from the specified distribution is added to each element of the one-hot input matrix.

#### Pretraining with data augmentations

Training with augmentations requires two main hyperparameters: first, a set of augmentations to sample from; and second, the maximum number of augmentations to be applied to a sequence. For each mini-batch during training, each sequence is randomly augmented independently. The number of augmentations to be applied to a given sequence has two possible settings in EvoAug: (1) `hard`, always equal to the maximum number of augmentations; or (2) `soft`, randomly select the number of augmentations for a sequence from 1 to the maximum number. Our experiments with Basset and DeepSTARR use the former setting, while our experiments with ChIP-seq datasets use the latter setting. Then, the subset of augmentations to be applied to the sequence is sampled randomly without replacement from the user-defined set of augmentations. After a subset of augmentations is chosen, the order in which multiple augmentations are applied to a single sequence is given by the following priority: inversion, deletion, translocation, insertion, reverse-complement, mutation, noise addition. Each augmentation is then applied stochastically for each sequence.

For the Basset and DeepSTARR models, each augmentation has an optimal setting that was determined from a hyperparameter search independently using the validation set (Supplementary Figs. 1, 3, and 4). For the Basset models, the hyperparameters were set to:

- mutation: `mutate_frac = 0.15`
- translocation: `shift_min = 0, shift_max = 30`
- insertion: `insert_min = 0, insert_max = 30`
- deletion: `delete_min = 0, delete_max = 30`
- reverse-complement: `rc_prob = 0.5`
- noise: `noise_mean = 0, noise_std` (standard deviation) = 0.3

For the DeepSTARR models, the hyperparameters were set to:

- mutation: `mutate_frac = 0.05`
- translocation: `shift_min = 0, shift_max = 20`
- insertion: `insert_min = 0, insert_max = 20`
- deletion: `delete_min = 0, delete_max = 30`
- reverse-complement: `rc_prob = 0`
- noise: `noise_mean = 0, noise_std =0.3`

When augmentations were used in combinations, the maximum number of augmentations was set to 3 for Basset and 2 for DeepSTARR. The same hyperparameter settings used in DeepSTARR analyses with all augmentations were used for the ChIP-seq analysis. For models trained with combinations of augmentations, the hyperparameters intrinsic to augmentations were set at the values identified above and the maximum number of augmentations per sequence was also determined through a hyperparameter sweep for each dataset (Supplementary Figs. 2 and 5).

Unless otherwise specified, all models were trained (with or without data augmentations) for 100 epochs using the Adam optimizer^35^ with an initial learning rate of 1 × 10^−3^ and a weight decay (*L*_2_ penalty) term of 1 × 10^−6^; additionally, we employed early stopping with a patience of 10 epochs and a learning rate decay that decreased the learning rate by a factor of 0.1 when the validation loss did not improve for 5 epochs. For each model trained, the version of the model with the highest-performing weights during its training, as measured by validation loss, is the version of the model whose performance is reported here.

#### Fine-tuning

Models that completed training with data augmentations were subsequently fine-tuned on the original dataset without augmentations. Fine-tuning employs the Adam optimizer with a learning rate of 1 × 10^−4^ and a weight decay (*L*_2_ penalty) term of 1 × 10^−6^ for 5 epochs. The model that yields the lowest validation loss was used for test time evaluation.

#### Evaluation

When evaluating models on validation or test sets, no data augmentations were used on input sequences. For models trained with an insertion augmentation (alone or in combination with other augmentations), each sequence is padded at the 3’ end with a stretch of random DNA of length `insert_max`.

### Interpretability Analysis

#### Filter interpretability

We visualized the first-layer filters of various Basset models according to activation-based alignments^36^ and compared how well they match motifs in the 2022 JASPAR nonredundant vertebrates database^37^ using Tomtom^38^, a motif search comparison tool. Matrix profiles MA1929.1 and MA0615.1 were excluded from filter matching to remove poor quality hits; low information content filters tend to have a high hit rate with these two matrix profiles. Hit rate is calculated by measuring how many filters matched to at least one JASPAR motif. Average q-value is calculated by taking the average of the smallest q-values for each filter among its matches.

#### Attribution analysis

SHAP-based^39^ attribution maps (implemented with `GradientShap` from the Captum package^40^) were used to generate sequence logos (visualized by Logomaker^41^) for sequences that exhibited high experimental enhancer activity for the Developmental promoter (i.e., task 0 in the DeepSTARR dataset). 1,000 random DNA sequences were synthesized to serve as references for each GradientShap-based attribution map. A gradient correction^42^ was applied to each attribution map. For comparison, this analysis was repeated for a DeepSTARR model that was trained without any augmentations and a fine-tuned DeepSTARR model that was pretrained with all augmentations (excluding inversions) with two augmentations per sequence.

### CAGI5 Challenge Analysis

The CAGI5 challenge dataset^21^ was used to benchmark model performance on variant effect predictions. This dataset contains massively parallel reporter assays (MPRAs) that measure the effect size of single-nucleotide variants through saturation mutagenesis of 15 different regulatory elements ranging from 187 nt to 600 nt in length. We extracted 600 nt sequences from the reference genome centered on each regulatory region of interest and converted it into a one-hot representation. Alternative alleles were then substituted correspondingly to construct the CAGI test sequences.

For a given Basset model, the output predictions of two input sequences, one with a centered reference allele and the other with an alternative allele, are made. The cell type-agnostic approach employed in this study uses the mean across these values to calculate a single scalar value, functional activity across cell types. The effect size is then calculated with the log-ratio of this single value for the alternative allele and reference allele, according to: log(alternative value / reference value).

To evaluate the variant effect prediction performance, Pearson correlation was calculated within each CAGI5 experiment between the experimentally measured and predicted effect sizes. The average of the Pearson correlation across all 15 experiments represents the overall performance of the model.

## Data and code availability

EvoAug Python package is deposited on the Python Package Index (PyPI) repository with documentation hosted on https://evoaug.readthedocs.io. The open-source project repository is available at GitHub, https://github.com/p-koo/evoaug. The code to reproduce analyses in this paper is available on GitHub, https://github.com/p-koo/evoaug_analysis. Data and model weights are available at Zenodo: doi.org/10.5281/zenodo.7265991 and doi.org/10.5281/zenodo.7277777.

## Acknowledgements

This work was supported in part by funding from the NIH grant R01HG012131 and the Simons Center for Quantitative Biology at Cold Spring Harbor Laboratory. This work was performed with assistance from the US National Institutes of Health Grant S10OD028632-01. The authors would also like to thank Jakub Kaczmarzyk for help setting up the *Python Package Index* and code documentation on ReadTheDocs.org.

## Author contributions statement

NKL and PKK conceived of the method, designed the experiments, and wrote the majority of the code base. NKL, ST, and ZT conducted experiments and analyzed the data. PKK oversaw the project. All authors interpreted the results and contributed to the manuscript.

---

**Supplementary Table 1.**
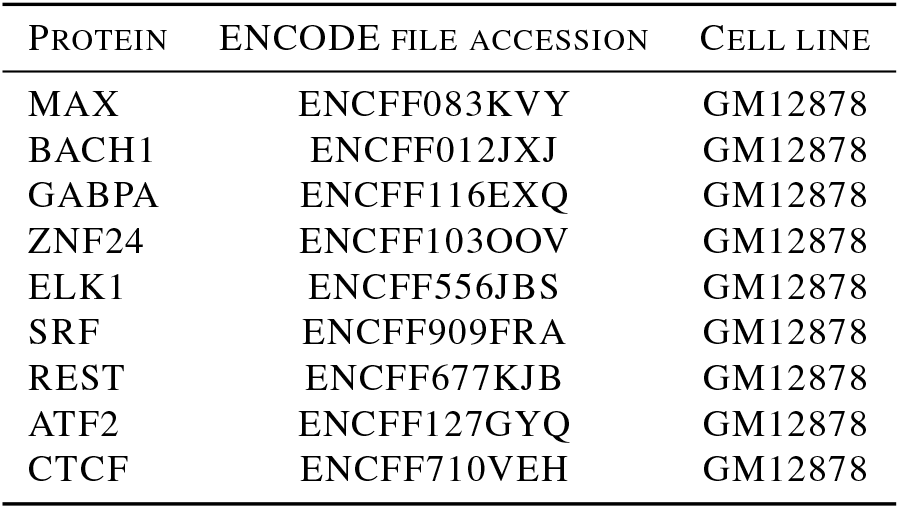
ENCODE ChIP-seq details. Ten representative TF ChIP-seq experiments in GM12878 cell line and a DNase-seq experiment (File accession: ENCFF235KUD) for the same cell line were downloaded from ENCODE. Table shows ENCODE file accession codes for all transcription factor proteins.

**Supplementary Table 2.** Computational cost for Basset models. Table shows the computational cost of EvoAug augmentations for Basset models, including the number of epochs, time per epoch, and the total training time, on a single NVIDIA RTX 2080ti GPU. The values represent the average across 5 independent trials and errors represent standard deviation of the mean.

| AUGMENTATION | EPOCHS | TIME PER EPOCH (S) | TOTAL TIME (MIN) |
| --- | --- | --- | --- |
| STANDARD | 14.0 ± 1.3 | 74 | 17.3 ± 1.6 |
| NOISE | 35.4 ± 9.7 | 98 | 57.8 ± 15.8 |
| RC | 25.4 ± 5.8 | 114 | 48.3 ± 11.0 |
| MUTATION | 52.4 ± 12.5 | 171 | 149.3 ± 35.625 |
| TRANSLOCATION | 25.6 ± 3.3 | 125 | 53.3 ± 6.9 |
| DELETION | 32.2 ± 6.1 | 172 | 92.3 ± 17.5 |
| INSERTION | 34.0 ± 8.3 | 174 | 98.6 ± 24.1 |
| NOISE+RC+INS | 87.6 ± 12.2 | 219 | 319.7 ± 44.5 |
| ALL | 85.4 ± 15.1 | 259 | 368.6 ± 65.2 |

**Supplementary Table 3.** Computational cost for DeepSTARR models. Table shows the computational cost of EvoAug augmentations for DeepSTARR models, including the number of epochs, time per epoch, and the total training time, on on a single NVIDIA RTX 2080ti GPU. The values represent the average across 5 independent trials and errors represent standard deviation of the mean.

| AUGMENTATION | EPOCHS | TIME PER EPOCH (S) | TOTAL TIME (MIN) |
| --- | --- | --- | --- |
| STANDARD | 22.4 ± 1.4 | 62 | 23.1 ± 1.4 |
| NOISE | 45.0 ± 11.1 | 76 | 57.0 ± 14.1 |
| MUTATION | 49.6 ± 8.0 | 138 | 114.1 ± 18.4 |
| TRANSLOCATION | 58.6 ± 12.4 | 108 | 105.5.1 ± 22.3 |
| DELETION | 56.6 ± 11.2 | 149 | 140.6 ± 27.8 |
| INSERTION | 52.8 ± 13.6 | 156 | 137.3 ± 35.4 |
| NOISE+INS | 81.8 ± 16.0 | 217 | 295.8 ± 57.9 |
| ALL | 74.4 ± 10.0 | 264 | 327.4 ± 44.0 |

**Supplementary Figure 1.**
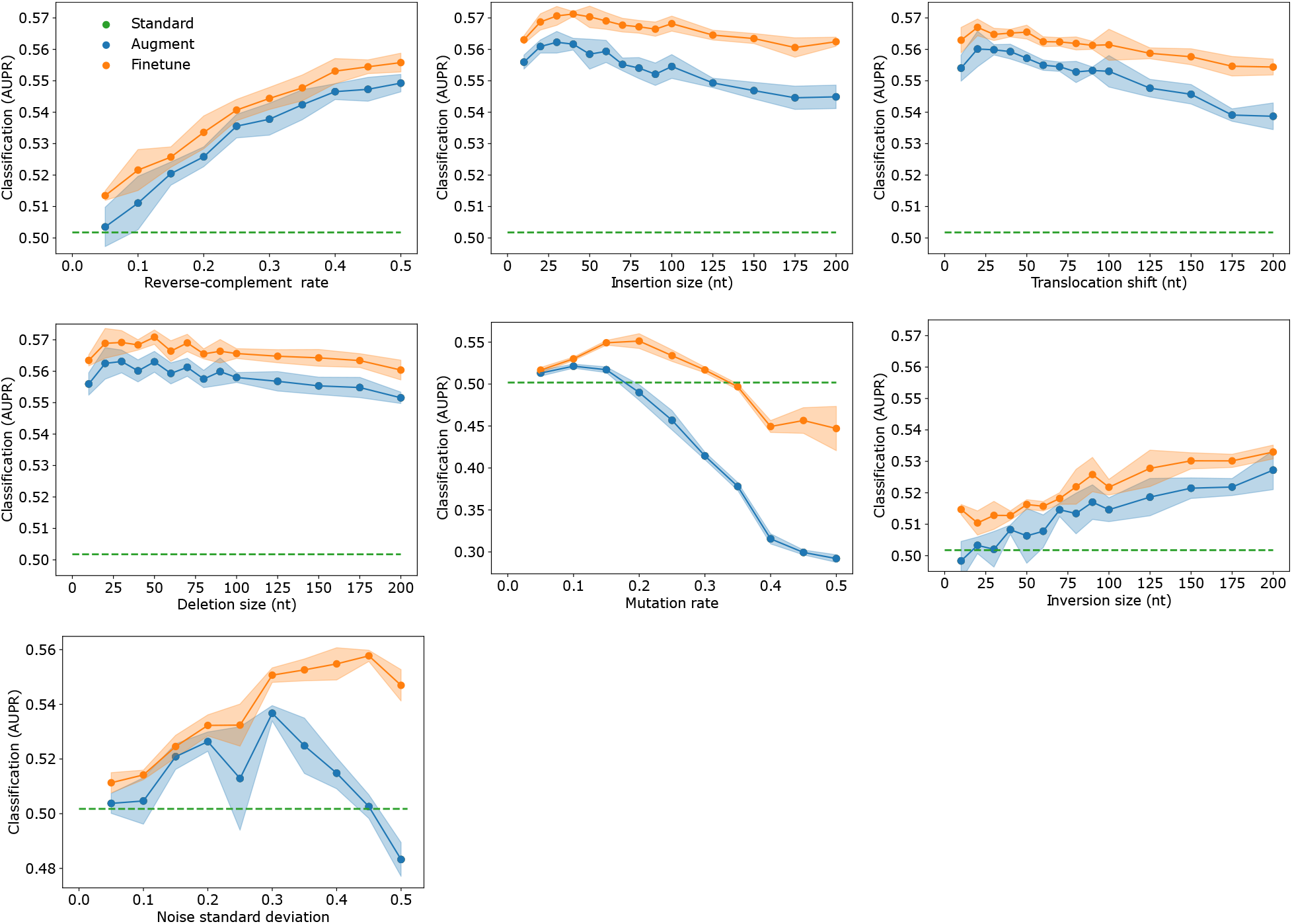
Hyperparameter sweep of each augmentation method for Basset. Each plot shows the average classification performance for different hyperparameter values intrinsic to each data augmentation method. Shaded region represents the standard deviation of the mean. Dashed line represents the performance without augmentations. Values reported are with n =5 trials with random initializations.

**Supplementary Figure 2.**
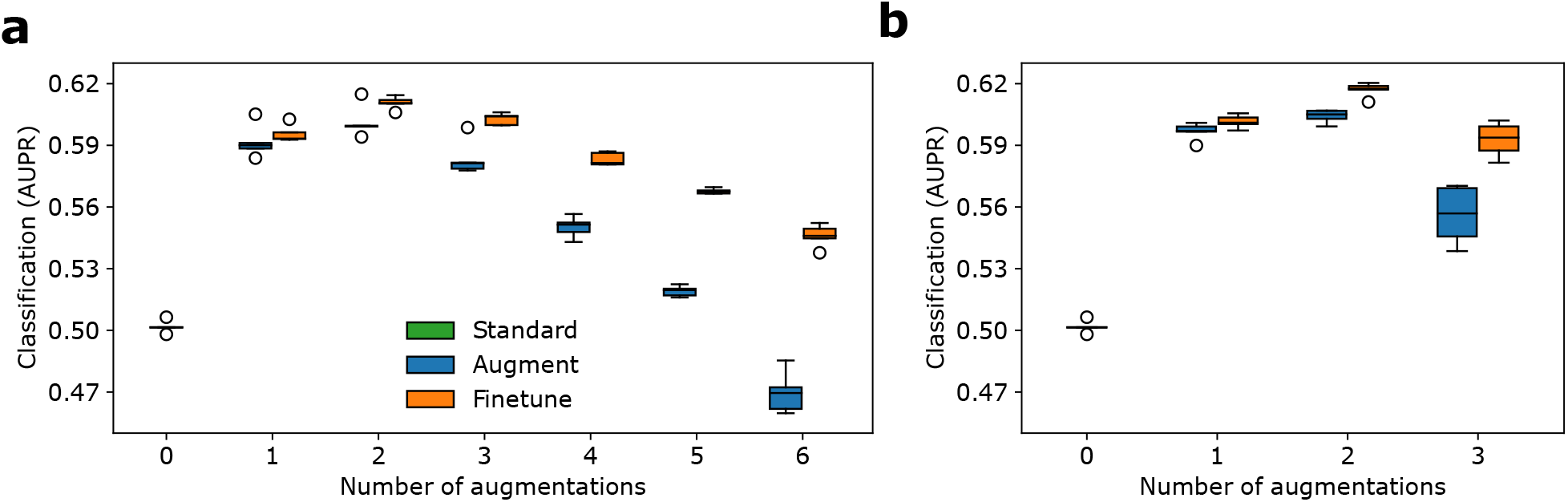
Sweep in number of applied augmentations for Basset. (**a**) Box-plot of classification performance for Basset models pre-trained with two combination strategies of (**a**) all and (**b**) insertion + translocation + deletion, and fine-tuned on original data. All represents reverse-complement, Gaussian noise, insertion, deletion, translocation, and mutation. Standard represents no augmentations during training. Each number in the *x*-axis represents the number of augmentations that are applied to each sequence in combinations during training. Box plots show the first and third quartiles, central line is the median, and the whiskers show the range of data with outliers removed. Values reported are with *n* = 5 trials with random initializations.

**Supplementary Figure 3.**
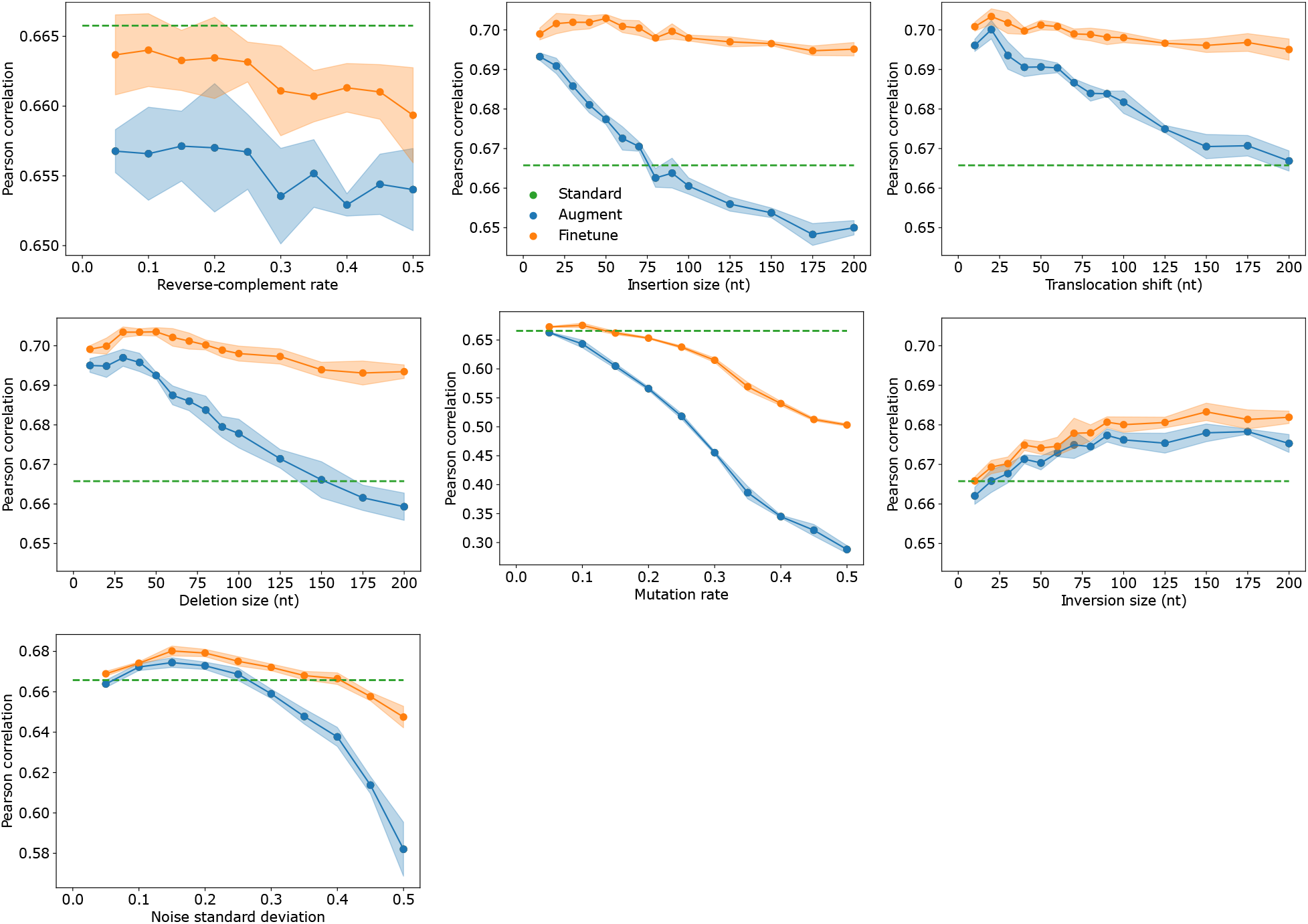
Hyperparameter sweep of each augmentation method for DeepSTARR, Developmental. Each plot shows the average classification performance for different hyperparameter values intrinsic to each data augmentation method. Shaded region represents the standard deviation of the mean. Dashed line represents the performance without augmentations. Values reported are with n =5 trials with random initializations.

**Supplementary Figure 4.**
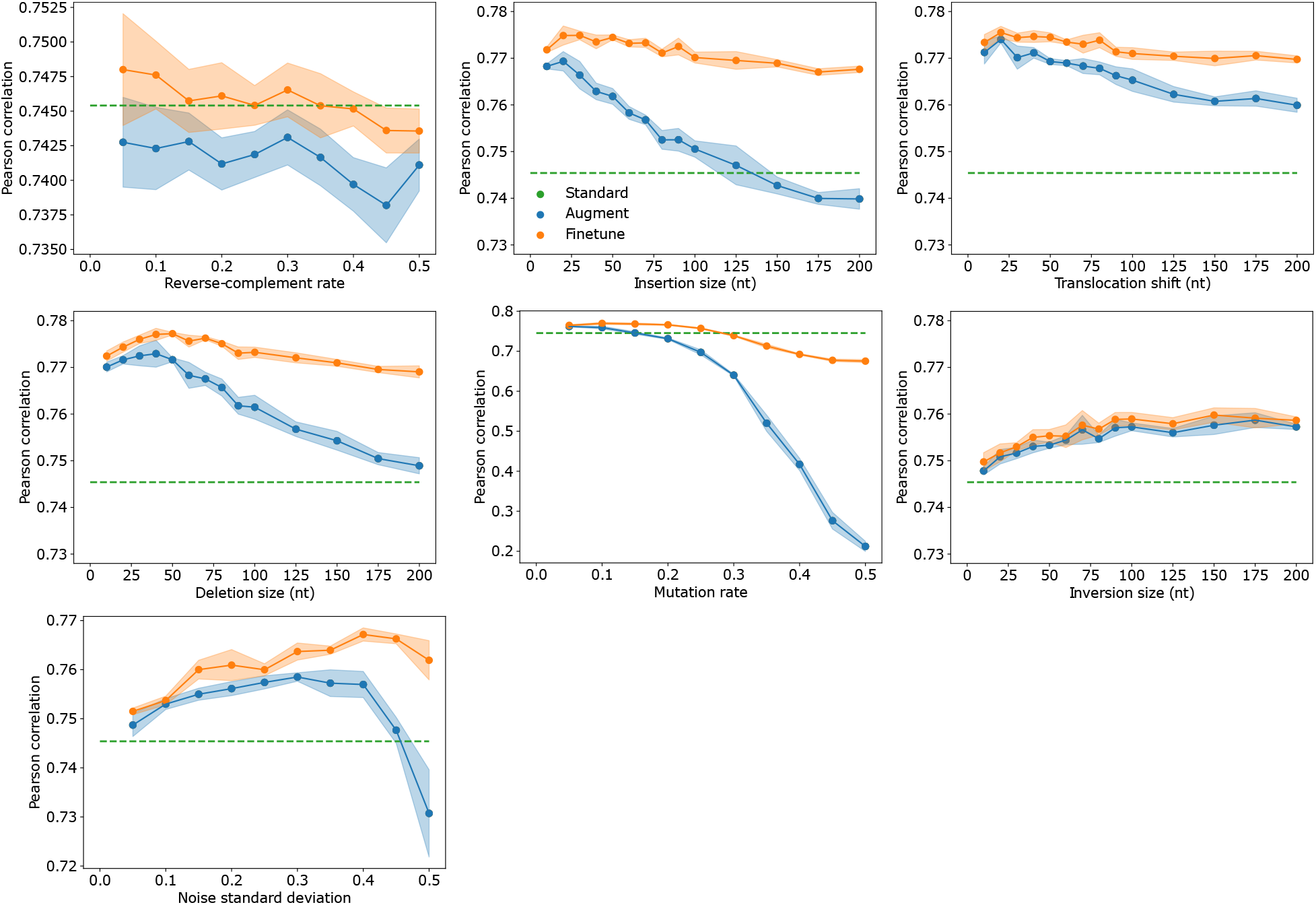
Hyperparameter sweep of each augmentation method for DeepSTARR, Housekeeping. Each plot shows the average classification performance for different hyperparameter values intrinsic to each data augmentation method. Shaded region represents the standard deviation of the mean. Dashed line represents the performance without augmentations. Values reported are with *n* = 5 trials with random initializations.

**Supplementary Figure 5.**
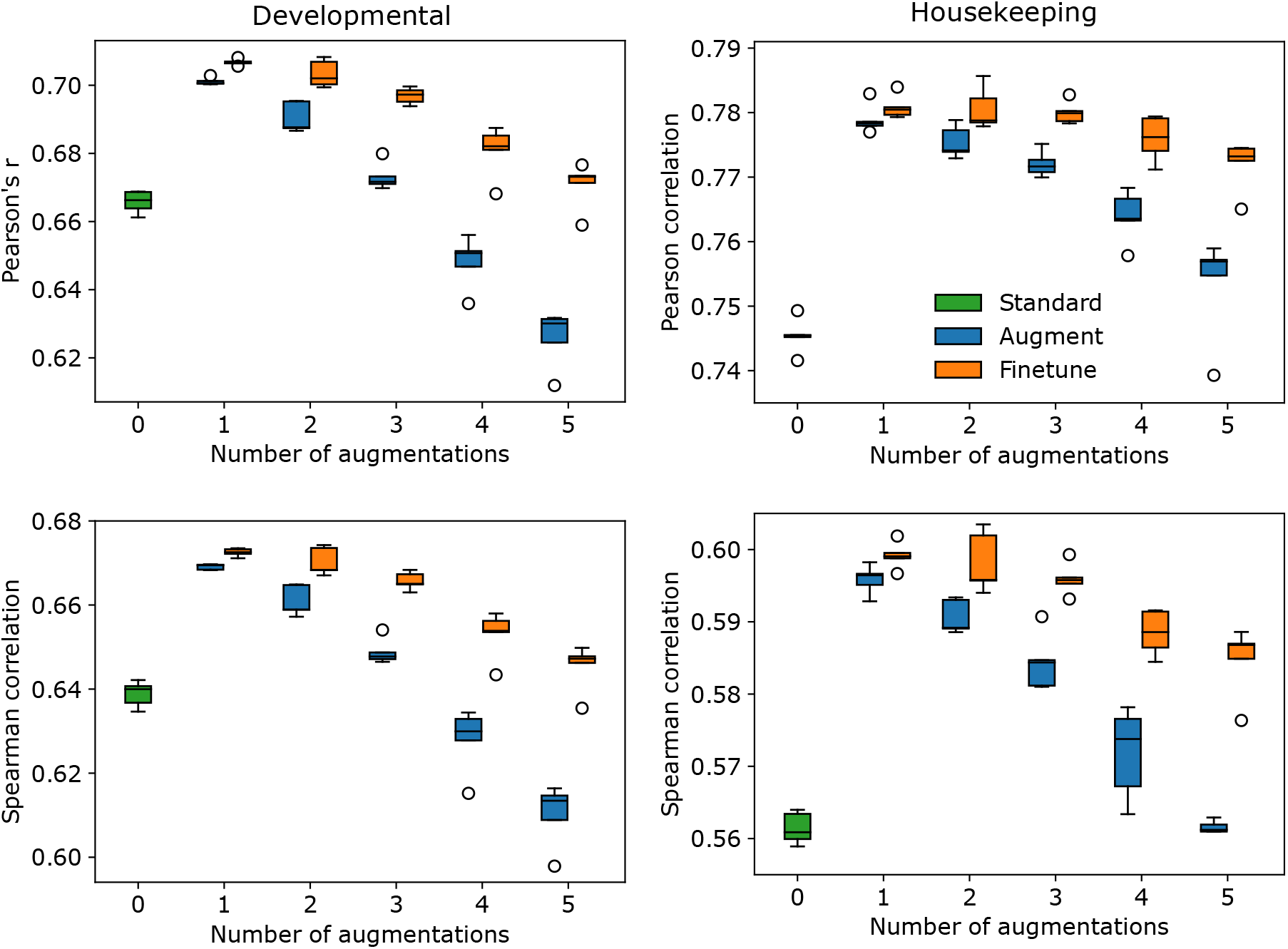
Sweep in number of applied augmentations for DeepSTARR. Box-plot of regression performance (Pearson’s *r* for top row and Spearman correlation for bottom row) for DeepSTARR models pre-trained with all augmentations—namely, reverse-complement, Gaussian noise, insertion, deletion, translocation, and mutation—for developmental (left) and housekeeping (right) conditions. Standard represents no augmentations during training. Each number in the x-axis represents the number of augmentations that are applied to each sequence in combinations during training. Box plots show the first and third quartiles, central line is the median, and the whiskers show the range of data with outliers removed. Values reported are with n =5 trials with random initializations.

**Supplementary Figure 6.**
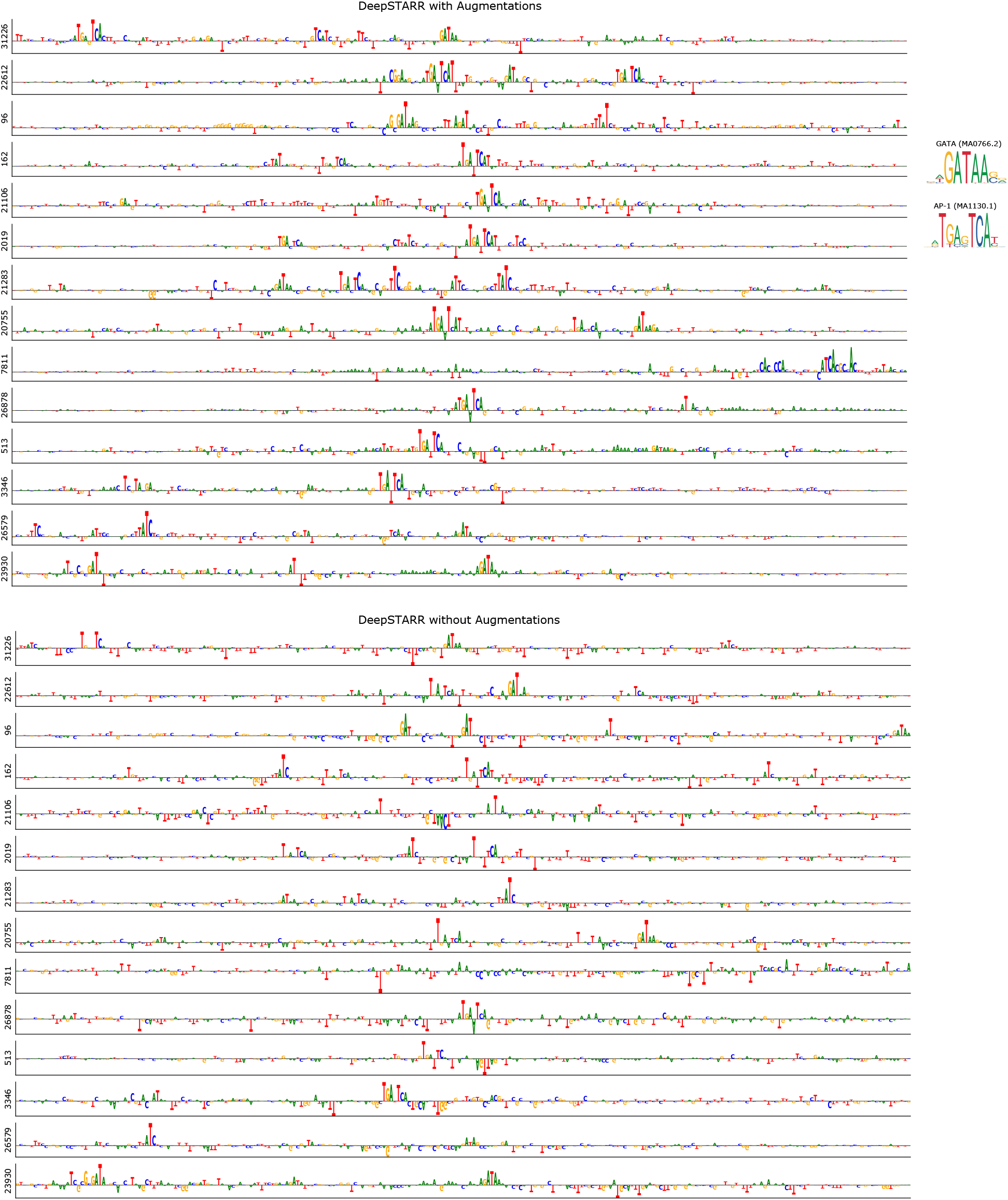
Attribution map comparison for DeepSTARR. Sequence logos of SHAP-based attribution maps for a fine-tuned DeepSTARR model that was pretrained with a combination of all augmentations up to 2 augmentations total per sequence (top) and DeepSTARR trained without any augmentations (bottom). The number on the y-axis represents the index of the sequence from the DeepSTARR test set. Sequence logos of GATA and AP1 are shown for visual comparison.

**Supplementary Figure 7.**
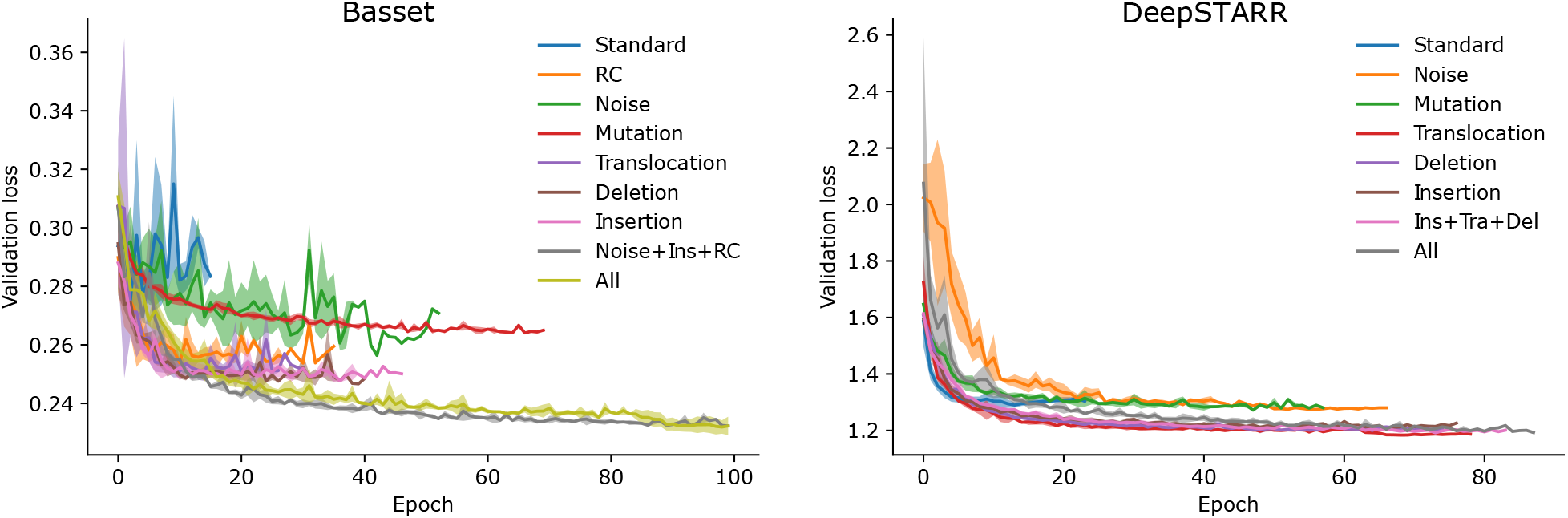
Training performance. The average validation loss at each epoch of training for various augmentations applied to Basset (left) and DeepSTARR (right). Values reported are with *n* = 5 trials with random initializations. Shaded region represents the standard deviation. Due to the different training times for each trial, standard deviation was calculated for epochs that contained at least 3 values.

